# HIC1 links retinoic acid signalling to group 3 innate lymphoid cell-dependent regulation of intestinal immunity and homeostasis

**DOI:** 10.1101/108241

**Authors:** Kyle Burrows, Frann Antignano, Alistair Chenery, Vladimir Korinek, T. Michael Underhill, Colby Zaph

## Abstract

The intestinal immune system must be able to respond to a wide variety of infectious organisms while maintaining tolerance to non-pathogenic microbes and food antigens. The Vitamin A metabolite retinoic acid (RA) has been implicated in the regulation of this balance, partially by regulating innate lymphoid cell (ILC) responses in the intestine. However, the molecular mechanisms of RA-dependent intestinal immunity and homeostasis remain elusive. Here we define a role for the transcriptional repressor Hypermethylated in cancer 1 (HIC1, ZBTB29) in the regulation of ILC responses in the intestine. Intestinal ILCs express HIC1 in a vitamin A-dependent manner. In the absence of HIC1, group 3 ILCs (ILC3s) are lost, resulting in increased susceptibility to infection with the bacterial pathogen *Citrobacter rodentium*. In addition, the loss of ILC3s leads to a local and systemic increase in IFN-γ-producing T cells that prevents the development of protective immunity against infection with the parasitic helminth *Trichuris muris*. Thus, RA-dependent expression of HIC1 in ILC3s regulates intestinal homeostasis and protective immunity.

**Author Summary:** Innate lymphoid cells (ILCs) are emerging as important regulators of immune responses at barrier sites such as the intestine. However, the molecular mechanisms that control this are not well described. In the intestine, the Vitamin A metabolite retinoic acid (RA) has been shown to be an important component of the homeostatic mechanisms. In this manuscript, we show that the RA-dependent transcription factor Hypermethylated in cancer 1 (HIC1, ZBTB29) is required for ILC homeostasis and function in the steady state as well as following infection with the bacterial pathogen *Citrobacter rodentium* or the helminth parasite *Trichuris muris*. Thus, HIC1 links RA signalling to intestinal immune responses. Further, our results identify HIC1 as a potential target to modulate ILC responses in vivo in health and disease.

## INTRODUCTION

The intestinal immune system is held in a tightly regulated balance between immune activation in response to potential pathogens and the maintenance of tolerance to innocuous antigens, such as food and commensal flora. Disruption of this balance can lead to the development of serious inflammatory disorders, such as food allergy or inflammatory bowel disease (IBD). A complex network of different immune cell types including dendritic cells (DCs), macrophages, innate lymphoid cells (ILCs), and T cells, are essential for both the induction of active immunity and the maintenance of intestinal homeostasis.

The vitamin A metabolite all-*trans*-retinoic acid (atRA) plays an important role in shaping intestinal immunity by regulating both the innate and adaptive immune systems. atRA that is generated by the metabolism of Vitamin A by intestinal epithelial cells (IECs) and a subset of CD103-expressing intestinal dendritic cells (CD103^+^ DCs) has been shown to directly affect the localization and function of lymphocytes. For example, atRA has been shown to induce expression of chemokine receptors (CCR9) and integrins (α4 and β7) that are associated with homing to, and retention in, the intestinal microenvironment (Iwata et al., 2004; Kim et al., 2015; Mora and von Andrian, 2009). In addition, atRA has been shown to control the balance of regulatory T (T_reg_) cells and CD4^+^ T helper 17 (T_H_17) cells in the intestine by promoting T_reg_ cell differentiation and inhibiting T_H_17 cell development (Sun et al., 2007; Mucida et al., 2007; Elias et al., 2008; Hill et al., 2008; Xiao et al., 2008; Takaki et al., 2008). Similarly, atRA controls the development of ILC subsets in the intestine, as mice raised on a Vitamin A-deficient (VAD) diet display reduced numbers of ILC3s (Spencer et al., 2014; Wilhelm et al., 2016), with one study showing a concomitant increase in ILC2 numbers and enhanced type 2 immunity within the intestine (Spencer et al., 2014). In addition, intestinal DC differentiation is influenced by atRA as mice raised on a VAD diet display reduced numbers of CD103^+^ CD11b^+^ DCs (Zeng et al., 2016). Thus, atRA-dependent processes are central to the function of intestinal T_H_ cells, ILCs, and DCs in vivo. However, the molecular mechanisms downstream of atRA signaling that control immune cell function and homeostasis remain unknown.

Hypermethylated in cancer 1 (HIC1, ZBTB29) is a transcriptional factor that was first identified as a gene that is epigenetically silenced in a variety of human cancers (Chen et al., 2003; Wales et al., 1995). HIC1 has been shown to regulate cellular proliferation, survival and quiescence in multiple normal and tumour cell lines (Lin et al., 2013; Valenta et al., 2006; Van Rechem et al., 2010; Chen et al., 2005). HIC1 is a member of the POZ and Kruppel/Zinc Finger and BTB (POK/ZBTB) family of transcription factors that consists of regulators of gene expression that are critical in a variety of biological processes (Kelly and Daniel, 2006). Importantly, several members of the POK/ZBTB family are key regulators in immune cell differentiation and function, including: BCL6, PLZF and ThPOK (Lee and Maeda, 2012; Nurieva et al., 2009; Savage et al., 2008; Wang et al., 2008; Muroi et al., 2008). Recently, we identified HIC1 as an atRA responsive gene in intestinal T_H_ cells and demonstrated a T cell-intrinsic role for HIC1 in the regulation of intestinal homeostasis as well as in development of several models of intestinal inflammation (Burrows, et al. 2017).

In this study, we show that HIC1 is critical for intestinal immune homeostasis by regulating ILC3s. Deletion of HIC1 in hemoatopoietic cells or RORγt-expressing cells results in a significant reduction in the number of ILC3s in the intestine. Further, we identify an ILC3-intrinic role for HIC1 in regulating intestinal T_H_ cell responses to commensal bacteria and immunity to infection with the intestinal pathogens *Citrobacter rodentium* and *Trichuris muris*. These results identify a central role for atRA-dependent expression of HIC1 in ILC3s in the regulation of intestinal immune responses.

## RESULTS

### HIC1 is expressed by intestinal ILCs and is critical for intestinal immune homeostasis

We have previously shown that HIC1 was highly expressed by T cells, DCs and macrophages within the intestinal lamina propria (LP) and intraepithelial compartments. (Burrows et al., 2017). Using mice with a fluorescent reporter gene inserted in the *Hic1* locus (*Hic1*^Citrine^ mice) (Pospichalova et al., 2011) we were able to determine that in addition to the previously identified populations, lineage-negative (lin^neg^) CD90.2^+^ CD127^+^ ILCs isolated from the intestinal lamina propria express HIC1 (**Fig 1A**). Similar to our previous results, HIC1 expression in ILCs was dependent on the availability of atRA, as *Hic1*^Citrine^ mice reared on a VAD diet did not express HIC1 in ILCs within the LP (**Fig 1B**). To determine the role of HIC1 in ILCs, we first crossed mice with *loxP* sites flanking the *Hic1* gene (*Hic1^fl/fl^* mice) with mice that express the Cre recombinase under control of the *Vav* promoter (*Vav*-Cre mice) to generate mice with a hematopoietic specific deletion of *Hic1* (*Hic1^Vav^* mice). Upon hematopoietic cell-specific deletion of HIC1 we observed a significant change in ILC populations in the LP. In the steady state, we observed significantly fewer RORγt^+^ ILCs (ILC3s) in the LP of *Hic1^Vav^* mice, with a significant reduction in the number of RORγt^+^ TBET+ ILC3s (**Fig 1C,D**). We also detected a small but significant increase in the number of CD4^+^ ILC3s (also known as lymphoid tissue inducer (LTi) cells) but no change in numbers of the canonical Gata3^+^ ILC (ILC2) population (**Fig 1C,D**). Thus HIC1 expression in hematopoietic cells is critical for regulation of ILC populations in the LP.

**Figure 1.**
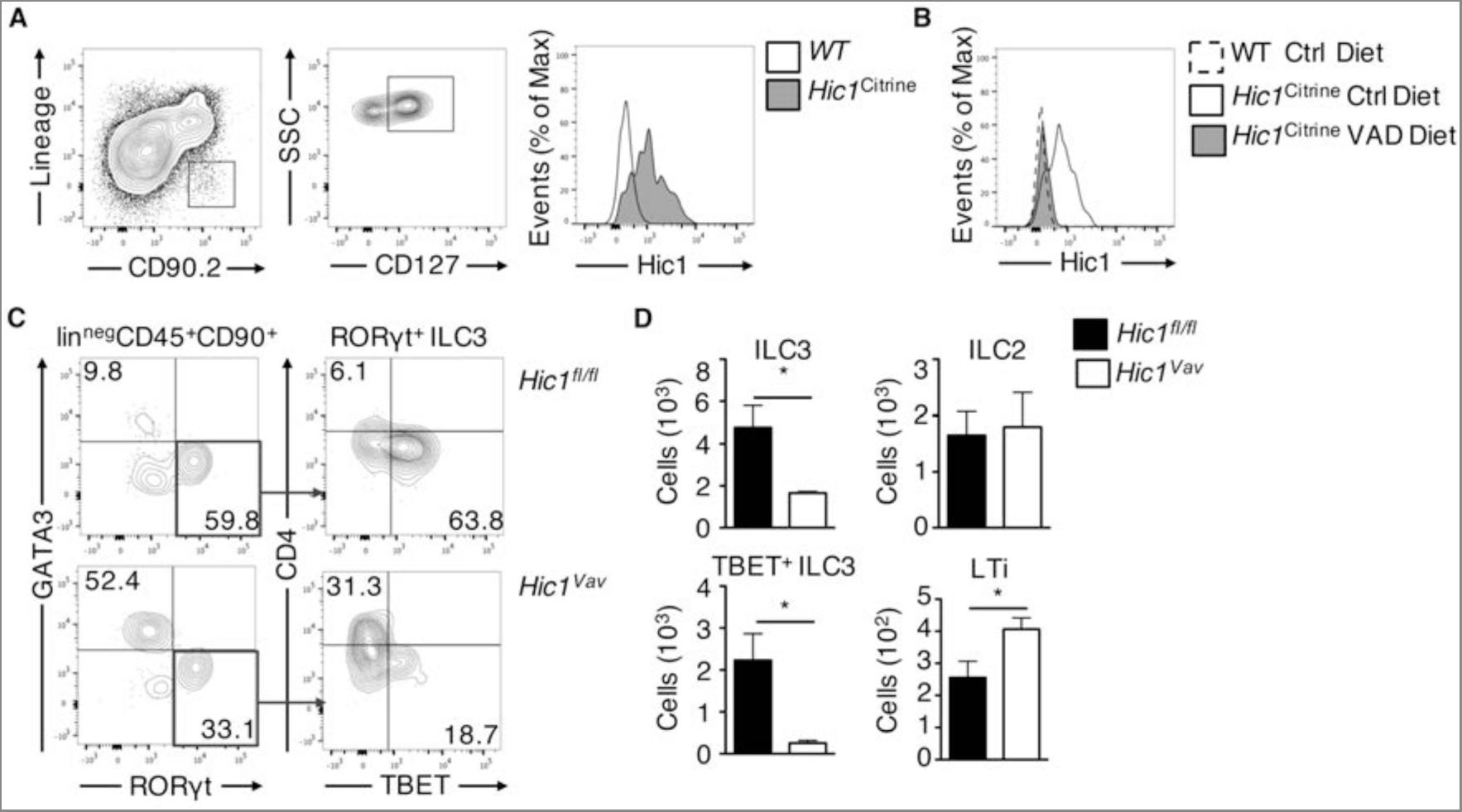
HIC1 is expressed by intestinal ILCs and is required for intestinal immune homeostasis. (A) ILCs (lin^neg^ CD90^+^ CD127^+^ cells) were analysed by flow cytometry for *Hic1^Citrine^* reporter expression from the intestinal lamina propria (LP). Data representative of 2 independent experiments (B) HIC1 reporter expression in intestinal LP ILCs from *Hic1^Citrine^* mice fed a control diet, *Hic1^Citrine^* mice fed a vitamin A deficient (VAD) diet, and controls fed a control diet was analysed by flow cytometry. Data are representative of 2 independent experiments. (C, D) Intestinal LPs from *Hic1^fl/fl^* and *Hic1^Vav^* mice at steady state were analysed by flow cytometry to enumerate populations of RORγt^+^ ILC3s, Gata3^+^ ILC2s, CD4^+^ ILC3s (LTis) and Tbet^+^ RORγt^+^ ILC3s. Data pooled from 2 independent experiments (n=4 per group). *, P < 0.05; Student’s *t* test. Errors bars indicate SEM.

### HIC1 does not regulate ILC precursors in the bone marrow

As we observed a significant reduction of ILC3s in the LP in the absence of HIC1, we next tested directly whether the lack of HIC1 affected the upstream development of ILC precursors in the bone marrow. ILCs develop in the bone marrow through a lineage pathway that begins with a common lymphoid progenitor (CLP) and progresses through an α4β7-expressing lymphoid progenitor (αLP), a common progenitor to all helper-like ILCs (ChILP) and, in the case of ILC2s, an ILC2 precursor (ILC2p) (Zook and Kee, 2016). Analysis of surface marker expression on lineage-negative, CD45^+^ bone marrow cells showed that HIC1 was not required for the development of CLP, αLP, ChILP or ILC2p populations (**Fig 2A,B**). Thus, the reduced number of ILC3s in the LP is not due to a reduced frequency of ILC precursors and suggests that HIC1 is required for ILC3 homeostasis in the periphery.

**Figure 2.**
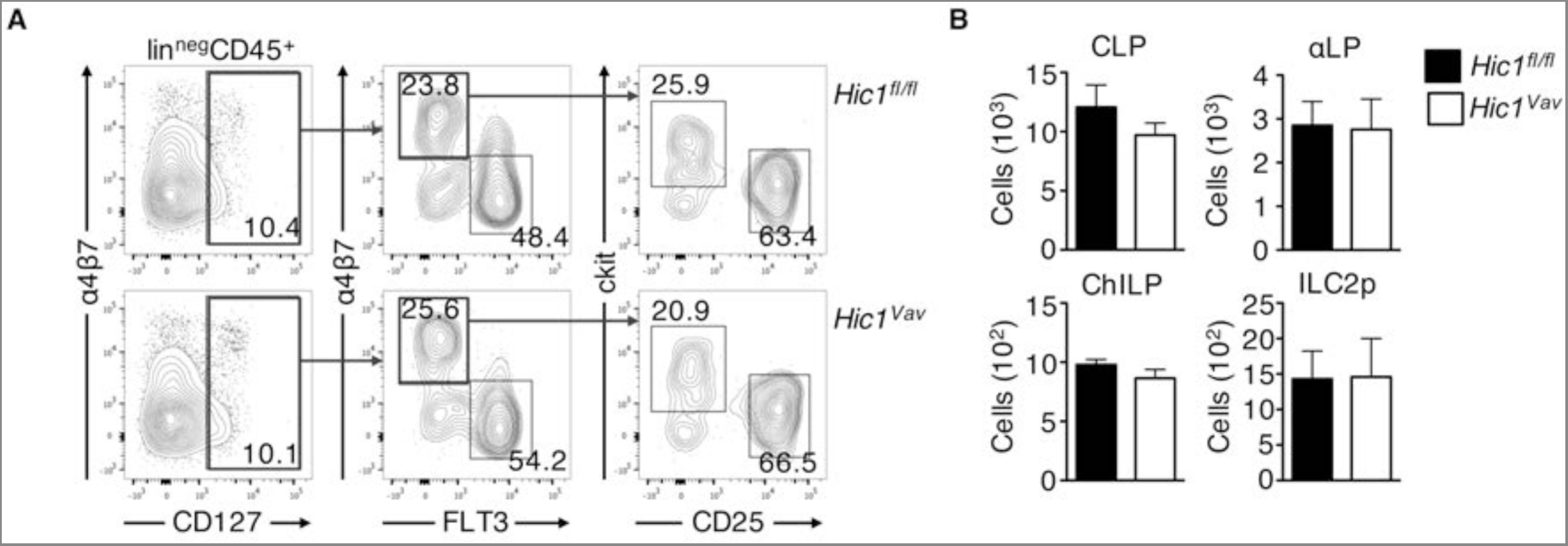
HIC1 does not regulate ILC precursors in the bone marrow. (A) Gating strategy and (B) cell numbers of CLPs (CD45^+^ lin^neg^ CD127^+^ Flt3^+^ α4β7^−^), α4β7^+^ lymphoid progenitors (αLP; CD45^+^ lin^neg^ CD127^+^ Flt3^−^ α4β7^+^), ChILPs (CD45^+^ lin^neg^ CD127^+^ Flt3^−^ α4β7^+^ CD25^−^ c-Kit^+^) and ILC2 progenitors (ILC2p; (CD45^+^ lin^neg^ CD127^+^ Flt3^−^ α4β7^+^ CD25^+^ c-Kit^−^) from bone marrow of *Hic1^Vav^* and *Hic1^fl/fl^* mice. Data are from two independent experiments (*n* = 4 per group). ns, not significant. *, P < 0.05; Student’s *t* test. Errors bars indicate SEM.

### Hematopoietic specific deletion of HIC1 results in susceptibility to intestinal bacterial infection

ILC3s have been shown to play a significant role in resistance to infection with the attaching and effacing bacterial pathogen *Citrobacter rodentium* (Satoh-Takayama et al., 2008; Sonnenberg et al., 2011). Following infection with *C. rodentium*, *Hic1^Vav^* mice exhibited enhanced weight loss and significantly higher bacterial burdens in the feces compared to *Hic1^fl/fl^* controls (**Fig 3A,B**). Furthermore, infected *Hic1^Vav^* mice but not *Hic1^fl/fl^* mice had dissemination of bacteria to the liver (**Fig 3C**), demonstrating a significant impairment in the intestinal barrier following infection. Associated with impaired bacterial containment and clearance were reduced levels of transcripts for the cytokines *Il17a* and *Il22*, as well as the intestinal antimicrobial peptide *Reg3g* (**Fig 3D**). Thus, expression of HIC1 within hematopoietic cells is critical to mount a proper immune response against *C. rodentium*.

**Figure 3.**
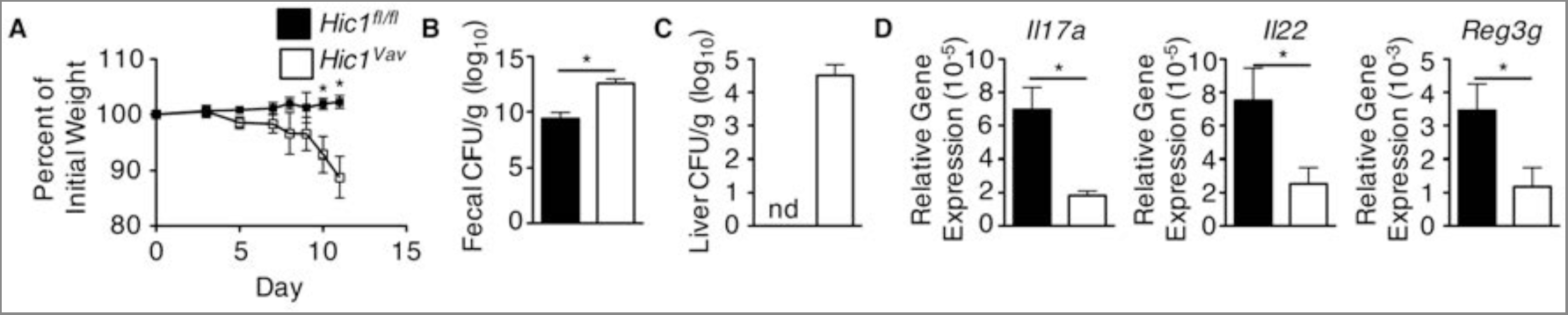
Hematopoietic deficiency of HIC1 results in susceptibility to *Citrobacter rodentium* infection. *Hic1^Vav^* and *Hic1^fl/fl^* mice were orally inoculated with *C.rodentium*. (A) Weight loss (percentage of initial weight) was calculated for each mouse over course of infection. (B, C) Bacterial loads (cfu/g) from fecal pellets (B) and liver (C) were measured at 11 days post inoculation. (D) Quantitative RT-PCR was performed to determine expression of *Il17a*, *Il22* and *Reg3g* from distal colon tissue 11 days post inoculation. Data are pooled from 2 independent experiments (n = 8-9 per group). *, P < 0.05; Student’s *t* test. Errors bars indicate SEM. nd, none detected.

### ILC3-intrinsic HIC1 expression is critical for defence against intestinal bacterial infection

As T cells, CD103^+^ CD11b^+^ DCs and ILC3s are all important in initiating and propagating ILC3/T_H_17 responses in the intestine (Persson et al., 2013; Kinnebrew et al., 2012; Collins et al., 2014; Klose and Artis, 2016) and these population are perturbed in *Hic1^Vav^* mice, we next sought to determine the effect of HIC1 deficiency in these specific cell populations during infection *C. rodentium*. We crossed *Hic1^fl/fl^* mice with mice expressing Cre under the control of either the *Cd4* promoter or *Itgax* promoter to generate T cell-specific (*Hic1^CD4^* mice) and dendritic cell-specific (*Hic1^CD11c^* mice) HIC1-deficient mice. Both *Hic^1CD4^* mice (**S1A-C Fig**) and *Hic1^CD11c^* mice (**S1D-F Fig**) were as resistant to infection with *C. rodentium* as control *Hic1^fl/fl^* mice, with equivalent weight loss, fecal bacterial burdens and expression of cytokines and antimicrobial peptide mRNA in the intestine. Thus, these results demonstrate that expression of HIC1 in T cells or CD11c-expressing cells is not required for immunity to bacterial infection and suggests loss of HIC1 in another cell population is responsible for the phenotype observed in *Hic1^Vav^* mice.

To determine the role of HIC1 expression in ILC3s during infection with *C. rodentium*, we crossed *Hic1^fl/fl^* mice with mice expressing Cre recombinase under the control of the Rorc promoter (*Hic1^Rorc^* mice). Following infection with *C. rodentium*, and similar to what we observed in the *Hic1^Vav^* mice, *Hic1^Rorc^* mice displayed increased weight loss, higher fecal bacterial burdens and increased bacterial dissemination than control *Hic1^fl/fl^* mice (**Fig 4A-C**). Associated with increased susceptibility was reduced expression of *IL17a*, *IL22* and *Reg3g* in intestinal tissues (**Fig 4D**). Based on our results, we hypothesized that HIC1 is an important regulator of ILC3 function during *C. rodentium* infection. To better examine the effect of HIC1 deletion on ILC3s, we examined the intestinal LP of *Hic1^fl/fl^* mice and *Hic1^Rorc^* mice at day 4 post *C. rodentium* infection. Enhanced susceptibility to infection with *C. rodentium* that we observed in Hic1Rorc mice correlated with the loss of ILC3s and reduced numbers IL-22-producing ILC3s in (**Fig 4E,F**). Taken together, these results suggest that expression of HIC1 in RORγt^+^ ILC3s is critical for resistance to intestinal bacterial infection.

**Figure 4.**
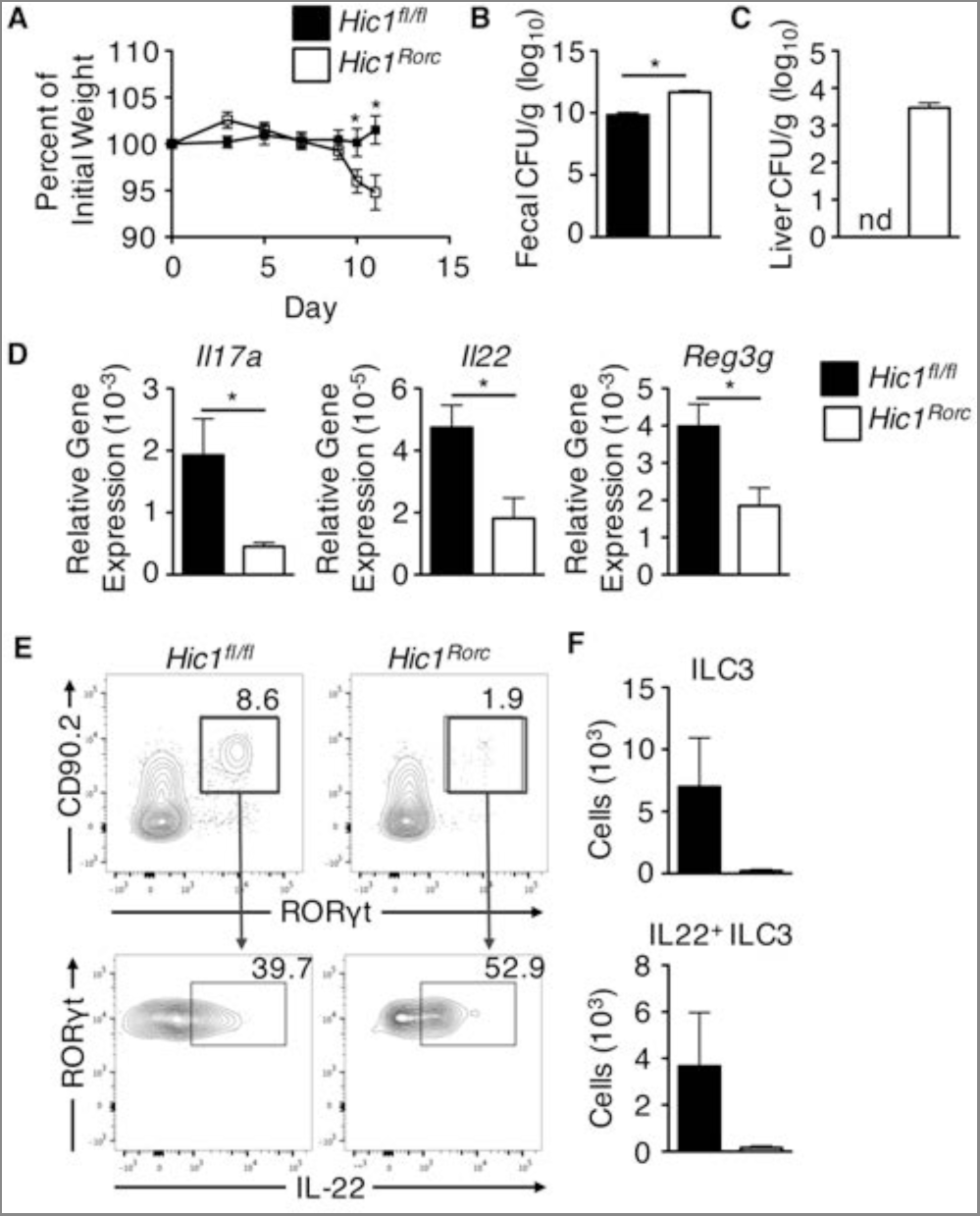
ILC3-intrinsic expression of HIC1 is required for immunity to *Citrobacter rodentium* infection. *Hic1^Rorc^* and *Hic1^fl/fl^* mice were orally inoculated with *C.rodentium*. (A) Weight loss (percentage of initial weight) was calculated for each mouse over course of infection. (b, c) Bacterial loads (cfu/g) from fecal pellets (B) and liver (C) were measured at 11 days post inoculation. (D) Quantitative RT-PCR was performed to determine expression of *Il17a*, *Il22* and *Reg3g* from distal colon tissue 11 days post inoculation. (E, F) Total ILC3s and IL-22 producing ILC3s from intestinal lamina propria were analysed by flow cytometry 4 days post inoculation. Data are pooled from two independent experiments (n = 7-8 per group). *, P < 0.05; Student’s *t* test. Errors bars indicate SEM. nd, none detected.

### Hematopoietic deficiency of HIC1 results in susceptibility to intestinal helminth infection

Our results show that HIC1 is an atRA-responsive factor that is critical for regulation of ILC3 homeostasis and function. As a recent study demonstrated that blockade of RA signalling results in dysfunctional ILC3 responses along with a compensatory increase in ILC2 responses and enhanced immunity to infection with the intestinal helminth parasite *Trichuris muris* (Spencer et al., 2014), we hypothesized that in addition to the defective ILC3 response observed in both *Hic1^Vav^* mice and *Hic1^Rorc^* mice, we would find enhanced ILC2 responses in the absence of HIC1. To test this, we infected *Hic1^fl/fl^* and *Hic1^Vav^* mice with *T. muris*. In contrast to our expectations, we did not observe a heightened protective ILC2/Th2 cell response, but found that *Hic1^Vav^* mice were susceptible to infection. *Hic1^Vav^* mice maintained a significant worm burden and enlarged mesenteric lymph node 21 days after infection (**Fig 5A,B**). Histological analysis revealed that *T. muris*-infected *Hic1^Vav^* mice displayed a reduced frequency of goblet cells, with parasites embedded within the caecal epithelium (**Fig 5C,D**). Associated with the increased susceptibility, *Hic1^Vav^* mice mounted a non-protective type 1 response following infection, as restimulation of the draining mesenteric lymph node (mLN) revealed a significant increase in secreted IFN-γ (**Fig 5E**) and expression of the *Ifng* in the intestinal tissues (**Fig 5F**), concomitant with a reduced type 2 response. We also observed a switch in the *T. muris*-specific antibody response from IL-4-dependent IgG1 to IFN-γ-dependent IgG2a in the serum of infected *Hic1^Vav^* mice (**Fig 5G**). Thus, loss of HIC1 in hematopoietic cells results in increased susceptibility to infection with *T. muris*.

**Figure 5.**
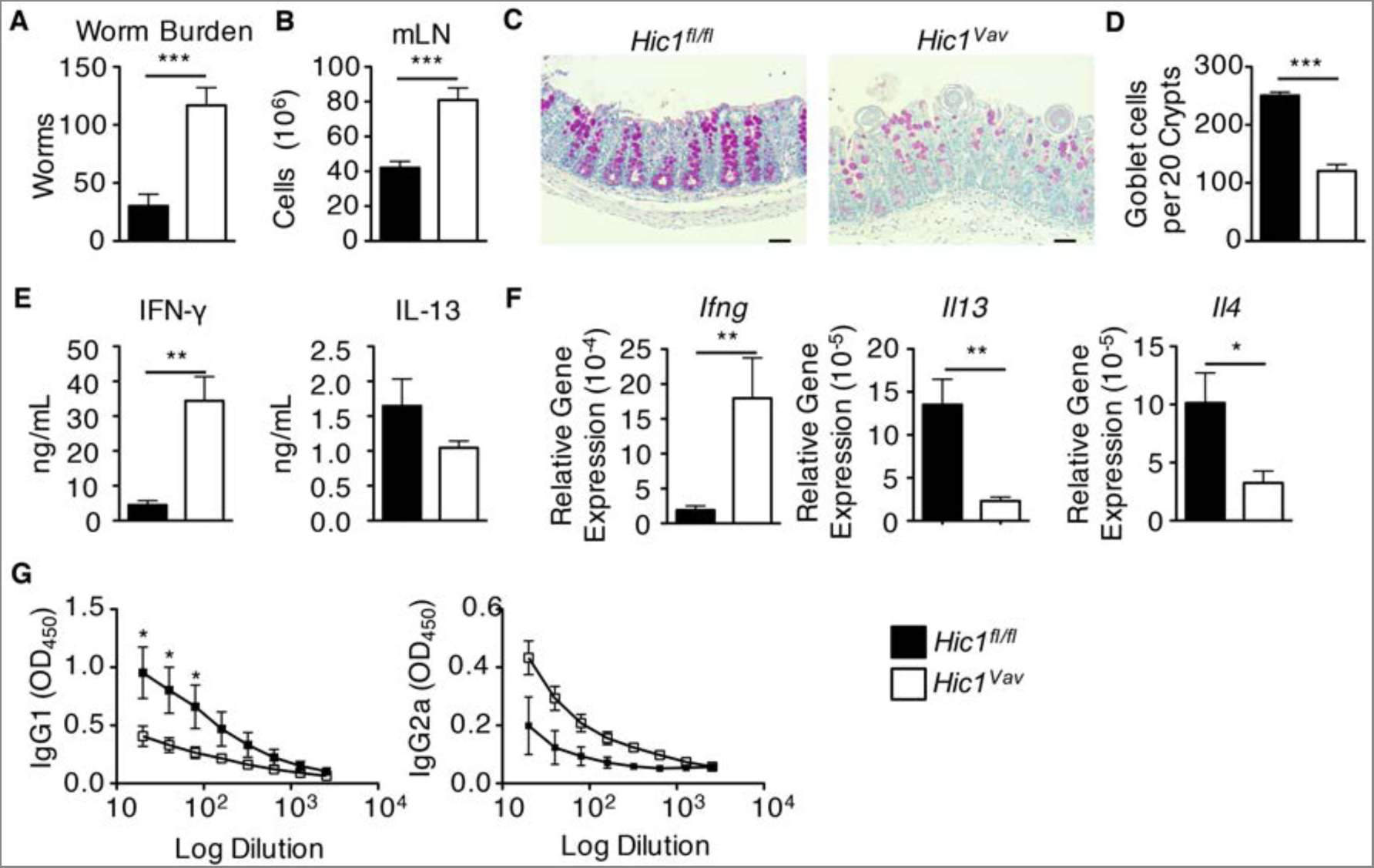
Hematopoieticexpression of HIC1 is required for resistance to *Trichuris muris* infection. *Hic1^Vav^* and *Hic1^fl/fl^* mice were infected with 200 *T. muris* eggs. (A) Worm burdens were determined microscopically from cecal contents 21 days post infection. (B) Total cells of mesenteric lymph node (mLN) were counted. (C) PAS-stained cecal sections, scale bar = 50 μm. (D) Goblet cells were quantified from PAS-stained cecal sections. (E) Supernatants of αCD3/CD28 restimulated mLN cells were evaluated for secretion of IFN-γ and IL-13 by ELISA. (F) Quantitative RT-PCR was performed to determine expression of *Il4*, *Il13*, and *Ifng* from proximal colon tissue. (G) Serum *Trichuris*-specific IgG1 and IgG2a levels were quantified by ELISA. (A–F) Data are pooled from three independent experiments (n=10-12 per group). (G) Data are representative of three independent experiments (n=4 per group). *, P < 0.05; **, P < 0.01; ***, P < 0.001; Student’s *t* test. Errors bars indicate SEM.

### HIC1 expression in T cells and dendritic cells is dispensable for immunity to *T. muris*

To determine which cell type required HIC1 expression to promote immunity to *T. muris*, we infected both *Hic1^CD4^* mice and *Hic1^CD11c^* mice. Strikingly, loss of HIC1 in T cells or CD11c-expressing cells had no effect on the development of protective immunity against *T. muris*. We observed equivalent expulsion of worms and expression of *Ifng* and *Il13* in the intestinal tissues between control *Hic1^fl/fl^* mice, *Hic1^CD4^* mice (**S2A,B Fig**) and *Hic1^CD11c^* mice (**S2C,D Fig**). Thus, we conclude that expression of HIC1 in T cells or CD11c-expressing cells is not required for immunity to *T. muris*.

### ILC3-intrinsic expression of HIC1 is required for immunity to *T. muris*

As there was no observable role for HIC1 in T cell or DC function during *T. muris* infection, we next sought to determine if there was a defect in HIC1 deficient ILC3s that prevent clearance of the parasite. Surprisingly, *Hic1^Rorc^* mice infected with *T. muris* were susceptible to infection, maintaining a significant parasite burden at day 21 post-infection (**Fig 6A**). Similar to *Hic1^Vav^* mice, we observed enlarged mesenteric lymph nodes (**Fig 6B**), as well as increased inflammatory cell infiltration, reduced number of goblet cells, submucosal edema, and parasites embedded within the caecal epithelium (**Fig 6C,D**). Consistent with the lack of protective immunity, we found that *Hic1^Rorc^* mice displayed high levels of secreted IFN-γ from restimulated mLN cells as well as increased expression of *Ifng* in intestinal tissues (**Fig 6E,F**), which is associated with increased levels of IFN-γ-dependent IgG2a antibodies in the serum (**Fig 6G**). However, we failed to observe reduced expression of type 2 cytokines in the mLN or intestine (**Fig 6E,F**), suggesting that the heightened levels of IFN-γ was promoting susceptibility in the context of a protective type 2 immune response. Thus, ILC3-intrinsic expression of HIC1 is critically required for the development of protective type 2 immune responses against *T. muris*.

**Figure 6.**
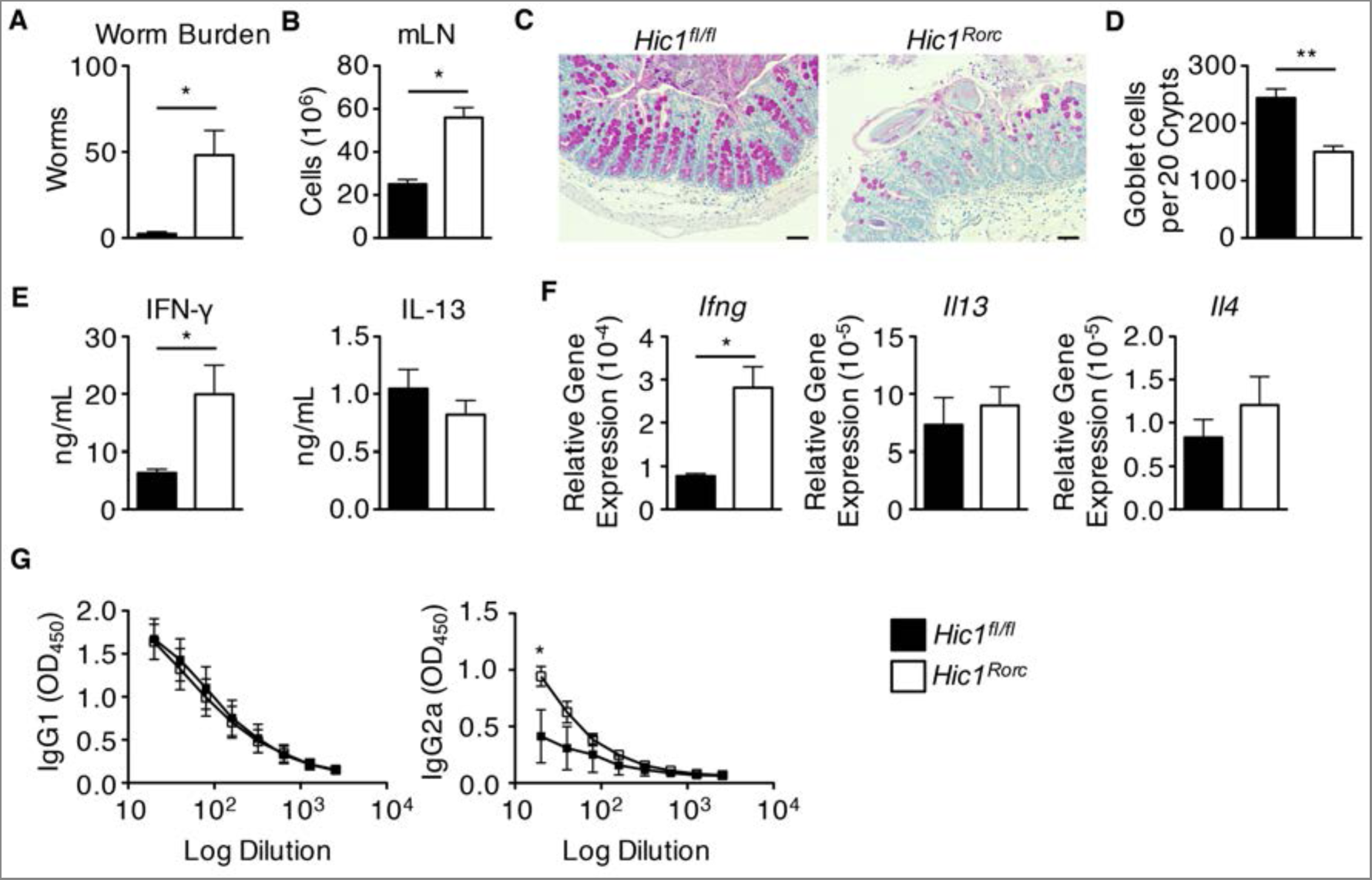
ILC3 specific deletion of HIC1 renders mice susceptible to *Trichuris muris* infection. *Hic1^Rorc^* and *Hic1^fl/fl^* mice were infected with 200 *T. muris* eggs. (A) Worm burdens were determined microscopically from cecal contents 21 days post infection. (B) Total cells of mesenteric lymph node (mLN) were counted. (C) PAS-stained cecal sections, scale bar = 50 μm. (D) Goblet cells were quantified from PAS-stained cecal sections. (E) Supernatants of αCD3/CD28 restimulated mLN cells were evaluated for secretion of IFN-γ and IL-13 by ELISA. (F) Quantitative RT-PCR was performed to determine expression of *Il4*, *Il13*, and *Ifng* from proximal colon tissue. (G) Serum *Trichuris*-specific IgG1 and IgG2a levels were quantified by ELISA. (A–F) Data are pooled from two independent experiments (n=8 per group). (G) Data are representative of two independent experiments (n=4 per group). *, P < 0.05; **, P < 0.01; ***, P < 0.001; Student’s *t* test. Errors bars indicate SEM.

### Neutralization of IFN-γ in *T. muris*-infected *Hic1^Rorc^* mice promotes resistance

Based on our results showing heightened levels of IFN-γ, we next asked whether antibody blockade of IFN-γ would render *Hic1^Rorc^* mice resistant to infection. We found that treatment of **T. muris**-infected *Hic1^Rorc^* mice with α-IFN-γ antibody promoted resistance, with a complete clearance of parasites in antibody-treated mice by day 21 post infection (**Fig 7A**). Increased resistance was associated with increased numbers of goblet cells (**Fig 7B,C**) along with reduced levels of IFN-γ and heightened levels of IL-4 and IL-13 (**Fig 7D,E**). Antibody treatment also resulted in reduced levels of *T. muris*-specific IgG2a and increased levels of IgG1 in the serum (**Figure 7F**). Thus, these results suggest that ILC3-specific expression of HIC1 is not required for resistance to *T. muris* infection when IFN-γ responses are neutralized.

**Figure 7.**
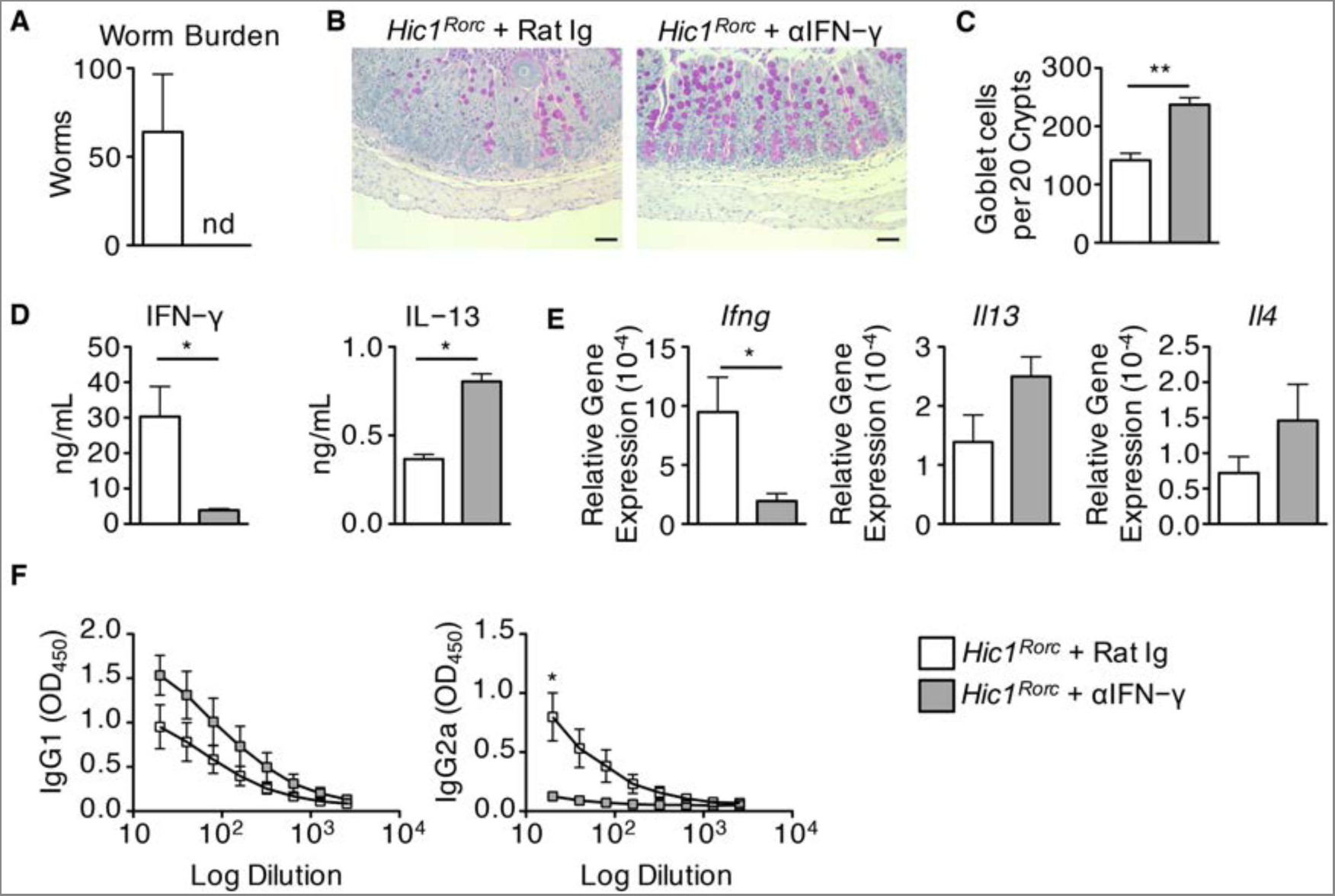
Neutralization of IFN_-γ_ in *Trichuris muris* - infected *Hic1^Rorc^* mice facilitates immunity to infection. *Hic1^Rorc^* mice were infected with 200 *T. muris* eggs and treated i.p. with antibodies against IFN-γ or an isotype control (Rat Ig). (A) Worm burdens were determined microscopically from cecal contents 21 days post infection. (B) PAS-stained cecal sections, scale bar = 50 μm. (C) Goblet cells were quantified from PAS-stained cecal sections. (D) Supernatants of αCD3/CD28 restimulated mLN cells were evaluated for secretion of IFN-*γ* and IL-13 by ELISA. (E) Quantitative RT-PCR was performed to determine expression of *Il4*, *Il13*, and *Ifng* from proximal colon tissue. (F) Serum Trichuris-specific IgG1 and IgG2a levels were quantified by ELISA. Data are from one experiment (n=4 per group). *, P < 0.05; **, P < 0.01; Student’s *t* test. Errors bars indicate SEM. nd, none detected.

### ILC3-intrinsic HIC1 is required to limit commensal bacteria specific T_H_ cell responses

ILC3s play a central role in intestinal immune homeostasis by limiting T cell responses against commensal bacteria (Hepworth et al., 2013; Hepworth et al., 2015). A subset of intestinal ILC3s can present commensal bacterial antigen to CD4^+^ T cells through MHCII but lack any co-stimulatory molecules and thus induce anergy in commensal-specific T cells (Hepworth et al., 2013). As we observed reduced numbers of ILC3s as well as enlarged mLNs and heightened levels of IFN-γ production from restimulated T cells from the mLN of naïve and *T. muris*-infected *Hic1^Rorc^* mice, we hypothesized that the increased IFN-γ production was due to dysregulated T cell responses to bacteria. Consistent with this, we observe a significant reduction in the number of regulatory MHCII^+^ ILC3s in the intestinal LP of *Hic1^Rorc^* mice at steady state (**Fig 8A,B**). Interestingly, analysis of CD4^+^ T cells from the mLN of *Hic1^Rorc^* mice at steady state reveals that in comparison to control mice, *Hic1^Rorc^* mice exhibited significantly increased frequencies of proliferating Ki67^+^ CD4^+^ T cells (**Fig 8D**), effector/effector memory CD44^high^ CD62L^low^ CD4^+^ T cells (**Fig 8E**) as well as IFN-γ^+^ CD4^+^ T cells (**Fig 8F**), indicative of disrupted immune cell homeostasis. Consistent with these responses being driven by commensal bacteria, oral administration of a cocktail of antibiotics to *Hic1^Rorc^* mice was associated with significantly reduced peripheral IFN-γ^+^ CD4^+^ T cells and CD44^high^ CD62L^low^ CD4^+^ T cells and mLN size (**Fig 8C-F**). Taken together, these results suggest that ILC3-intrinsic expression of HIC1 is required to limit commensal specific T_H_ cell responses in the steady state.

**Figure 8.**
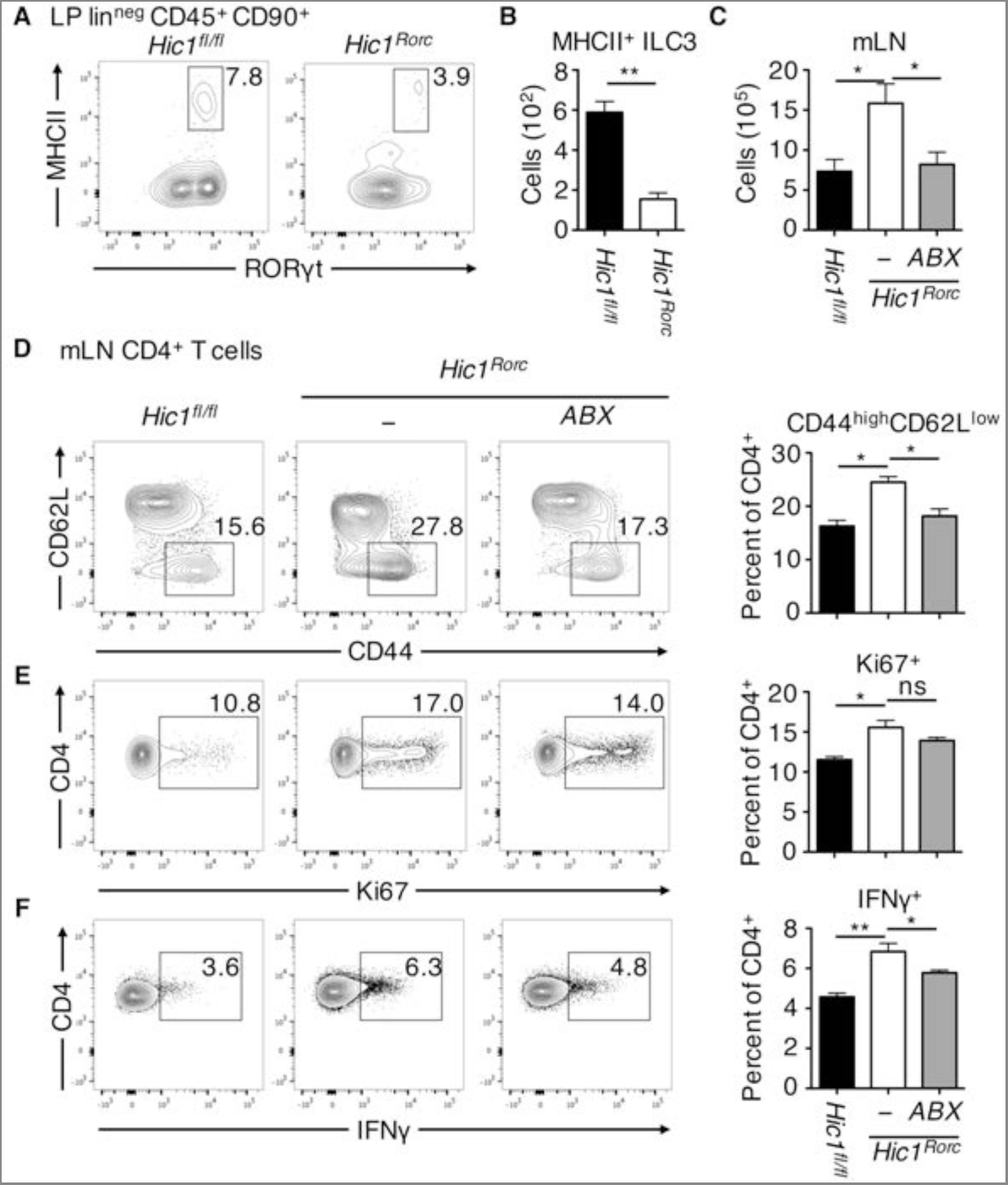
ILC3 expression of HIC1 is required to limit CD4^+^ T_H_ cell responses to commensal bacteria. (A, B) MHCII^+^ RORγt^+^ ILC3s from intestinal lamina propria of *Hic1^Rorc^* and *Hic1^fl/fl^* mice were analysed by flow cytometry (C–F) Mesenteric lymph node (mLN) from *Hic1^fl/fl^* and *Hic1^Rorc^* mice treated with or without antibiotics (Abx) in their drinking water were analyzed for total cell numbers (C), and analysed by flow cytometry for frequency of Ki67^+^ CD4^+^ T cells (D), frequency of CD44^high^ CD62L^low^ CD4^+^ T cells (E), and frequency of IFN-γ-producing CD4^+^ T cells (F). (A, B) Data are pooled from two independent experiments (n=4-5 per group). (C–F) Data are pooled from two independent experiments (n=5-7 per group). *, P < 0.05; **, P < 0.01; Student’s *t* test. Errors bars indicate SEM. ns, not significant.

## Discussion

Our results demonstrate that in the steady state, HIC1 is expressed by intestinal ILCs in a Vitamin A-dependent manner. In the absence of HIC1, we observed a dramatic decrease in intestinal ILC3 numbers, which was associated with a failure to clear *C. rodentium* infection. In addition, the reduction of regulatory MHCII^+^ ILC3s resulted in an increased frequency IFN-γ-producing T cells locally and systemically. The heightened levels of IFN-γ in the absence of HIC1 inhibited the ability of *Hic1^Vav^* mice and *Hic1^Rorc^* mice to mount a protective T_H_2 cell-associated immune response against *T. muris* infection. Together, these results highlight an important role for HIC1 not only in regulating intestinal immune homeostasis but also in mounting proper immune responses to diverse intestinal infections.

In the absence of HIC1, we found a significant reduction in the number of ILC3s with no effect on ILC2s in the intestine. Specifically, there were reduced numbers of RORγt^+^ TBET^+^ ILC3s and an increase in the number of CD4^+^ LTi cells. This is consistent with studies that have demonstrated that these two lineages have distinct developmental pathways; LTi cells would develop in the fetus while TBET^+^ ILC3s develop postnatally and rely on environmental signals (Klose et al., 2013; Sanos et al., 2009; Spencer et al., 2014). Interestingly, it has been shown that atRA signalling is also important for generation of LTi cells in the fetus (van de Pavert et al., 2014). However, our results suggest that HIC1 is not involved in fetal LTi formation, as we find no differences in LTi numbers or lymphoid structures in the absence of HIC1. Further, the development of ILC progenitor cells in the bone marrow is not perturbed by loss of HIC1, suggesting that the primary role of HIC1 is to regulate the development and function of adult cells in the periphery.

Resistance to intestinal infection with *C. rodentium* is mediated by IL-22, and ILC3s are the predominant IL-22-producing cell population during the first week of infection (Zheng et al., 2008; Sonnenberg et al., 2011). There are contradictory studies on which ILC3 populations are key for resistance to *C. rodentium* with both CD4^+^ LTis and natural cytotoxicity receptor (NCR)^+^ ILC3s each being described as either individually critical or redundant (Rankin et al., 2016; Sonnenberg et al., 2011; Song et al., 2015). Another study looking at TBET^+^ ILC3s (which include NCR^+^ ILC3s) demonstrated that TBET expression in a subset of ILC3s is critical for resistance to *C. rodentium* infection (Rankin et al., 2013). Our results are consistent with a role for NCR^+^ or TBET^+^ ILC3s in immunity to *C. rodentium* as *Hic1^CD4^* mice (deficient for HIC1 in T cells and LTi) are resistant to infection while *Hic1^Rorc^* mice (deficient for HIC1 in T cells and all ILC3s) are susceptible. Thus, HIC1 expression in ILC3s is critical for immunity to *C. rodentium*.

In addition to a reduction in ILC3s, a recent study identified that mice raised on a VAD diet displayed increased ILC2 numbers and heightened type 2 immunity to helminth infection in the intestine (Spencer et al., 2014). However, in the absence of HIC1, we did not observe an increase in ILC2 numbers nor increased resistance to infection with **T. muris**. Instead, we detected increased production of IFN-γ and an inability to mount a protective T_H_2 cell response to *T. muris* infection. Further, treatment of *T. muris*-infected *Hic1^Rorc^* mice with a neutralizing antibody against IFN-γ rendered the mice resistant to infection, demonstrating that HIC1-dependent responses are dispensable in the absence of IFN-γ and that the effects of RA on ILC2s are likely independent of HIC1.

Taken together, these results establish a role for the transcriptional repressor HIC1 as an atRA-responsive cell-intrinsic regulator of ILC3 cell function in the intestine, and identify a potential regulatory pathway that could be targeted to modulate ILC3 responses in the intestine.

## METHODS

### Ethics statement

Experiments were approved by the University of British Columbia Animal Care Committee (Protocol number A13-0010) and were in accordance with the Canadian Guidelines for Animal Research.

### Mice

The generation of *Hic1*^Citrine^ mice has been described (Pospichalova et al., 2011) and *Hic1^fl/fl^* mice will be described elsewhere (manuscript in preparation). *Cd4*-Cre mice were obtained from Taconic, *Vav*-Cre mice were obtained from T. Graf (Centre for Genomic Regulation, Barcelona, Spain) and *CD11c*-Cre (B6.Cg-Tg(Itgax-cre)1-1Reiz/J) and RORc-Cre (B6.FVB-Tg(RORc-cre)1Litt/J) mice were obtained from the Jackson Laboratory (Bar Harbor, ME, USA). Animals were maintained in a specific pathogen-free environment at the UBC Biomedical Research Centre animal facility.

### Diet Studies

Vitamin A-deficient (TD.09838) diet was purchased from Harlan Teklad Diets. At day 14.5 of gestation, pregnant females were administered the vitamin A-deficient diet and maintained on diet until weaning of litter. Upon weaning, females were returned to standard chow, whereas weanlings were maintained on special diet until use.

### Isolation of Lamina Propria Lymphocytes

Peyer’s patches were removed from the small intestine, which was cut open longitudinally, briefly washed with ice-cold PBS and cut into 1.5 cm pieces. Epithelium was stripped by incubated in 2mM EDTA PBS for 15 minutes at 37°C and extensively vortexed. Remaining tissue was digested with Collagenase/Dispase (Roche) (0.5 mg/mL) on a shaker at 250 rpm, 37°C, for 60 minutes, extensively vortexed and filtered through a 70µm cell strainer. The flow-through cell suspension was centrifuged at 1500rpm for 5 min. The cell pellet was re-suspend in 30% Percoll solution and centrifuged for 10 minutes at 1200 rpm. The pellet was collected and used as lamina propria lymphocytes.

### Antibodies and flow cytometry

Absolute numbers of cells were determined via hemocytometer or with latex beads for LP samples. Intracellular cytokine (IC) staining was performed by stimulating cells with phorbol 12-myristate 13-acetate (PMA), ionomycin, and Brefeldin-A (Sigma) for 4 hours and fixing/permeabilizing cells using the eBioscience IC buffer kit. All antibody dilutions and cell staining were done with PBS containing 2% FCS, 1 mM EDTA, and 0.05% sodium azide. Fixable Viability Dye eFluor 506 was purchased from eBioscience to exclude dead cells from analyses. Prior to staining, samples were Fc-blocked with buffer containing anti-CD16/32 (93, eBioscience) and 1% rat serum to prevent non-specific antibody binding. Cells were stained with fluorescent conjugated anti-CD11b (M1/70), anti-CD11c (N418), anti-CD19 (ID3), anti-CD5 (53-7.3), anti-CD8 (53.67), anti-CD3 (KT3)(2C11), anti-NK1.1 (PK136), anti-B220 (atRA-6B2), anti-Ter119 (Ter119), anti-Gr1 (RB6-8C5) produced in house, anti-CD4 (GK1.5), anti-CD25 (PC61.5), anti-CD45 (30-F11), anti-CD90.2 (53-2.1), anti-Gata3 (TWAJ), anti-RORgt (B2D), anti-Tbet (eBio4B10), anti-Flt3 (A2F10), anti-ckit (ACK2), anti-TCRβ (H57-597), anti-MHCII (I-A/I-E) (M5/114.15.2), anti-F4/80 (BM8), anti-α4β7 (DATK32), anti-IFN-γ (XMG1.2), anti-IL-22 (IL22JOP), anti-IL-13 (eBio13A), anti-Ki67 (SolA15) purchased from eBioscience, anti-CD127 (5B/199), anti-CD62L (MEL-14), anti-CD44 (IM7), anti-CD64 (X54.5/7.1.1) purchased from BD Biosciences. Data were acquired on an LSR II flow cytometer (BD Biosciences) and analysed with FlowJo software (TreeStar).

### *Citrobacter rodentium* infection

Mice were infected by oral gavage with 0.1 ml of an overnight culture of Luria-Bertani (LB) broth grown at 37°C with shaking (200 rpm) containing 2.5 × 10^8^ cfu of *C. rodentium* (strain DBS100) (provided by B. Vallance, University of British Columbia, Vancouver, British Columbia, Canada). Mice were monitored and weighed daily throughout the experiment and sacrificed at various time points. For enumeration of *C. rodentium*, fecal pellets or livers were collected in pre-weighed 2.0 ml microtubes containing 1.0 ml of PBS and a 5.0 mm steel bead (Qiagen). Tubes containing pellets or livers were weighed, and then homogenized in a TissueLyser (Retche) for a total of 6 mins at 20 Hz at room temperature. Homogenates were serially diluted in PBS and plated onto LB agar plates containing 100 mg/ml streptomycin, incubated overnight at 37°C, and bacterial colonies were enumerated the following day, normalizing them to the tissue or fecal pellet weight (per gram).

### *Trichuris muris* infection

Propagation of Trichuris muris eggs and infections were performed as previously described (Antignano et al., 2011). Mice were infected with approximately 150 – 200 embryonated *T. muris* eggs by oral gavage and monitered over a period of 21 days. Sacrificed mice were assessed for worm burdens by manually counting worms in the ceca using a dissecting microscope. Cecal tissues were fixed overnight in 10% buffered formalin and paraffin-embedded. A total of 5-µm-thick tissue sections were stained with periodic acid–Schiff (PAS) for histological analysis. mLNs were excised and passed through a 70 µm cell strainer to generate a single-cell suspension. mLN cells (4 × 10^6^/mL) were cultured for 72 h in media containing 1 µg/mL each of antibodies against CD3 (145-2C11) and CD28 (37.51; eBioscience, San Diego, CA). Cytokine production from cell-free supernatant was quantified by ELISA using commercially available antibodies (eBioscience).

### In vivo neutralization of IFN-γ

Mice were infected with *T. muris* as described above. On day 4 post infection, mice were injected i.p. with 500 µg of either control IgG or anti-IFN-γ (XMG1.2) (produced in-house by AbLabBiologics, UBC (Vancouver, BC)), constituted in sterile PBS. Mice were repeatedly injected thereafter on days 8, 12, and 16 prior to sacrifice on day 21.

### RNA isolation and quantitative real-time PCR

Tissues were mechanically homogenized and RNA was extracted using the TRIzol method according to the manufacturer's instructions (Ambion). cDNA was generated using High Capacity cDNA reverse transcription kits (Applied Biosystems). Quantitative PCR was performed using SYBR FAST (Kapa Biosystems) and SYBR green-optimized primer sets run on an ABI 7900 real-time PCR machine (Applied Biosystems). Cycle threshold (C_T_) values were normalized relative to beta-actin (*Actb*) gene expression. The primers used were synthesized de novo: *Il4* forward 5’- TCGGCATTTTGAACGAGGTC - 3’ and reverse 5’- CAAGCATGGAGTTTTCCCATG-3’; *Il13* forward 5’- CCTGGCTCTTGCTTGCCTT-3’ and reverse 5’- GGTCTTGTGTGATGTTGCTCA-3’; *Il17a* forward 5’-AGCAGCGATCATCCCTCAAAG-3’ and reverse 5’-TCACAGAGGGATATCTATCAGGGTC-3’; *Il22* forward 5’-ATGAGTTTTTCCCTTATGGGGAC-3’ and reverse 5’-GCTGGAAGTTGGACACCTCAA-3’; *Ifng* forward 5’-GGATGCATTCATGAGTATTGCC-3’ and reverse 5’-CCTTTTCCGCTTCCTGAGG-3’; *Reg3g* forward 5’-CCGTGCCTATGGCTCCTATTG- 3’ and reverse 5’-GCACAGACACAAGATGTCCTG −3’ *Actb* forward 5’-GGCTGTATTCCCCTCCATCG-3’ and reverse 5’-CCAGTTGGTAACAATGCCATGT-3’.

### Serum ELISA

Serum was collected from mice 21 days post-infection with *T.muris*. Immulon plates (Thermo Fischer Scientific, NY) were coated with 5 µg/mL of dialyzed *T. muris* antigen overnight at 4°C. Wash buffer was PBS containing 0.05% Tween 20. Plates were blocked and serum samples were diluted in 3% bovine serum albumin in PBS/0.05% Tween 20. Serum samples were incubated on plates for 1 hour at room temperature. Plates were then incubated with rat anti-mouse IgG1 or IgG2a conjugated to horseradishperoxidase (BD Pharmingen, CA) for 1 hour at room temperature. Plates were developed using 3,3’,5,5’- tetramethylbenzidine (TMB) substrate (Mandel Scientific, ON) and stopped with 1N HCl. Plates were read at 450 nm on a Spectramax 384 (Molecular Devices, CA).

## Antibiotic treatment

Mice received antibiotics (0.5 g/l of each ampicillin, gentamicin, neomycin and metronidazole, 0.25g/l vancomycin, with 4 g/l Splenda for taste) in their drinking water from weaning until euthanization (~10 week of age).

## Statistics

Data are presented as mean ± S.E.M. A two-tailed Student’s t-test using GraphPad Prism 5 software determined statistical significance. Results were considered statistically significant with P < 0.05.

## AUTHOR CONTRIBUTIONS

K.B., F.A., A.C., and C.Z. designed and performed the experiments. T.M.U. and V.K. provided assistance and contributed reagents and materials. K.B. and C.Z. wrote the manuscript.

## ACKNOWLEDGMENTS

We would like to thank R. Dhesi, L. Rollins (BRC core), A. Johnson (UBCFlow), M. Williams (UBC AbLab), T. Murakami (BRC Genotyping), I. Barta (BRC Histology), and all members of BRC mouse facility for excellent technical assistance. This work was supported by the Canadian Institutes of Health Research's (CIHR) Canadian Epigenetics, Environment and Health Research Consortium (grant 128090 to C. Zaph) and operating grants (MOP-89773 and MOP-106623 to C. Zaph) and an Australian National Health and Medical Research Council (NHMRC) project grants (APP 1104433 and APP1104466 to C. Zaph). F. Antignano is the recipient of a CIHR/Canadian Association of Gastroenterology/Crohn’s and Colitis Foundation of Canada postdoctoral fellowship. C. Zaph is a Michael Smith Foundation for Health Research Career Investigator and a Veski Innovation Fellow.

---

**Supplemental Figure 1.**
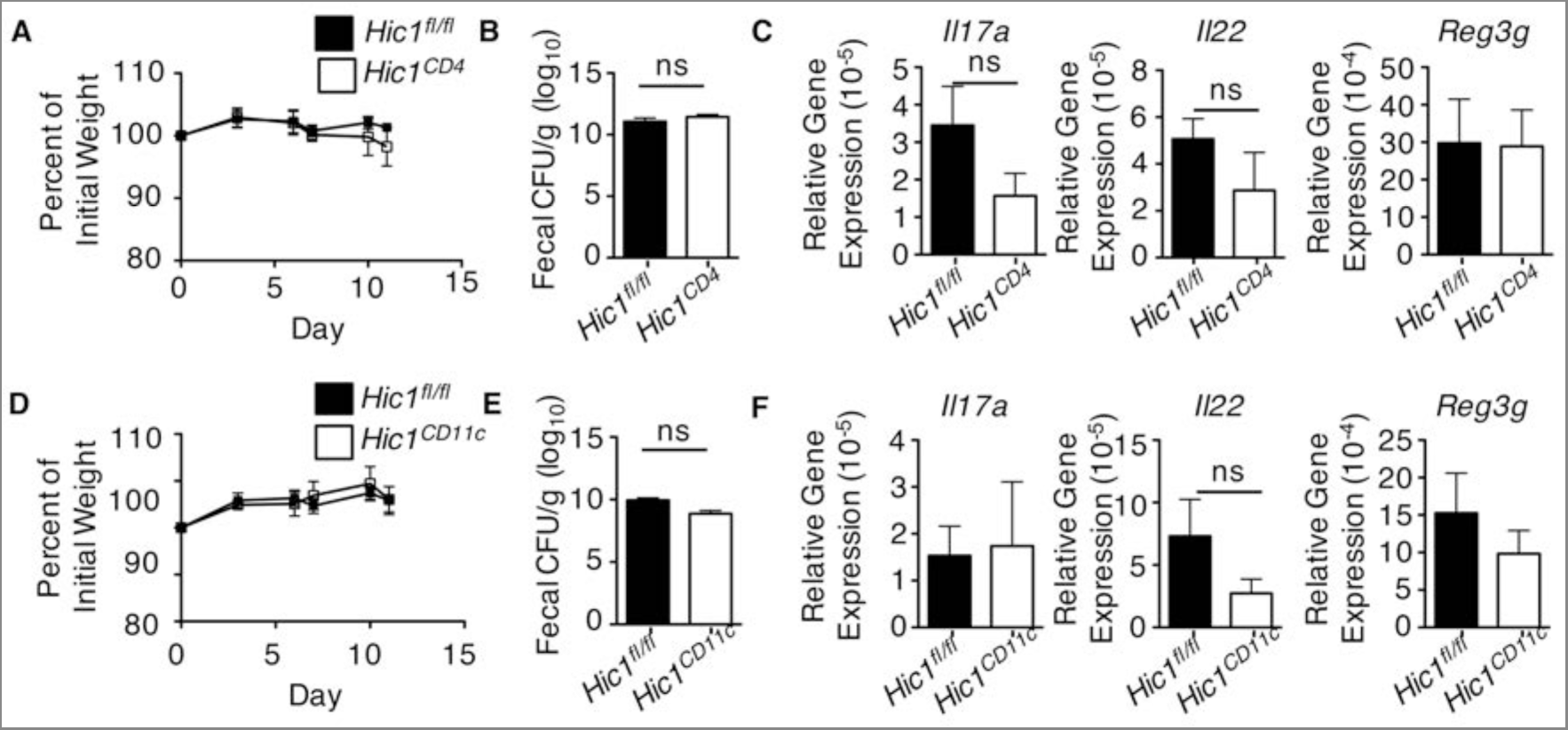
HIC1 expression in T cells and dendritic cells is not required for immunity to *Citrobacter rodentium* infection. (A–C) *Hic1^CD4^* and *Hic1^fl/fl^* mice or (D–f) *Hic1^CD11c^* and *Hic1^fl/fl^* mice were orally inoculated with *C. rodentium*. (A,D) Weight loss (percentage of initial weight) was calculated for each mouse over course of infection. (B,E) Bacterial loads (cfu/g) from fecal pellets were measured at 11 days post inoculation. (C,F) Quantitative RT-PCR was performed to determine expression of *Il17a*, *Il22* and *Reg3g* from distal colon tissue 11 days post inoculation. Data are pooled from 2 independent experiments (n = 6 per group, (A–C)) or (n = 5-6 per group, (D–F)). *, P < 0.05; Student’s *t* test. Errors bars indicate SEM. ns, not significant.

**Supplemental Figure 2.**
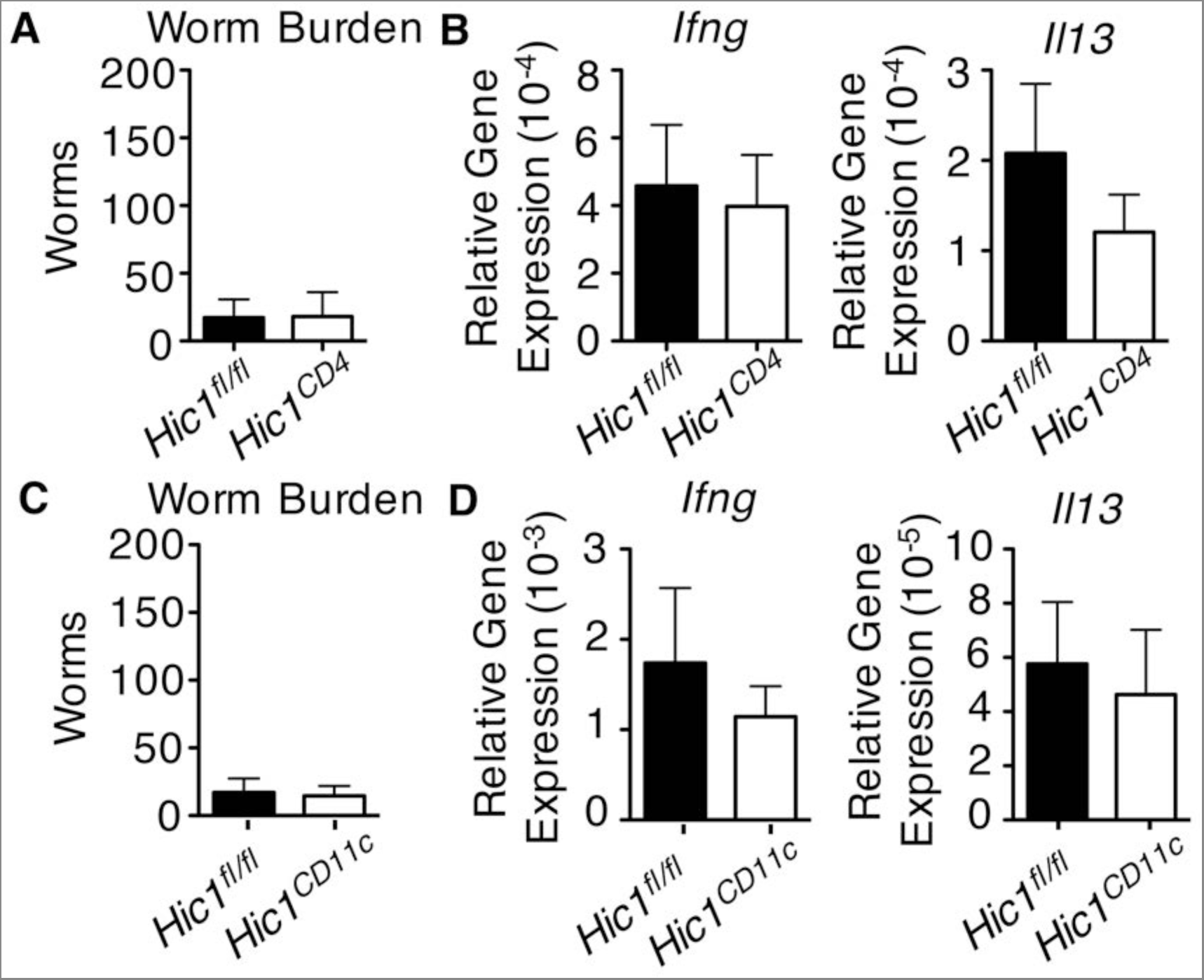
HIC1 expression in T cells and dendritic cells is dispensable for immunity to *T. muris*. (A,B) *Hic1^CD4^* and *Hic1^fl/fl^* mice or (C,D) *Hic1^CD11c^* and *Hic1^fl/fl^* mice were infected with 200 *T. muris* eggs. (A,C) Worm burdens were determined microscopically from cecal contents 21 days post infection. (B,D) Quantitative RT-PCR was performed to determine expression of *Il4*, *Il13*, and *Ifng* from proximal colon tissue. Data are pooled from 2 independent experiments (n = 8 per group). *, P < 0.05; Student’s *t* test. Errors bars indicate SEM. ns, not significant.

